# Protective Anti-Fibrotic Effect of Liraglutide and Pirfenidone Combination Therapy on Liver Fibrosis in Rats: Effects on Autophagy and NLRP3 Inflammasome

**DOI:** 10.1101/2025.08.19.671010

**Authors:** Zeynab Yousefi, Rayan Rajabi, Saeed Karima, Abbas Sahebghadam Lotfi, Mitra Nourbakhsh

**Affiliations:** Metabolic Disorders Research Center, Endocrinology and Metabolism Molecular-Cellular Sciences Institute, Tehran University of Medical Sciences, Tehran, Iran; Basic and Molecular Epidemiology of Gastrointestinal Disorders Research Center, Research institute for Gastroenterology and Liver Diseases, Shahid Beheshti University of Medical Sciences, Tehran, Iran; Oncopathology Research Center, Iran University of Medical Sciences, Tehran, Iran; Department of Clinical Biochemistry, School of Medicine, Shahid Beheshti University of Medical Sciences (SBMU), Tehran, Iran; Department of Clinical Biochemistry, Faculty of Medical Sciences, Tarbiat Modares University, Tehran, Iran; Department of Clinical Biochemistry, School of Medicine, Iran University of Medical Sciences, Tehran, Iran

**Author notes:** Corresponding Author: Abbas Sahebghadam Lotfi, Professor of clinical Biochemistry, Department of Clinical Biochemistry, Faculty of Medical Sciences, Tarbiat Modares University., Address: Tehran, Jalal Ale Ahmad, Nasr, P.O. Box: 14115-111. Co-Corresponding Author: Mitra Nourbakhsh, Professor of clinical Biochemistry, Department of Clinical Biochemistry, School of Medicine, Iran University of Medical Sciences, Tehran, Iran, Address: Tehran, Hemmat Highwaay, Iran University of Medical Sciences, P.O. Box: 1419614535.

**Keywords:** Pirfenidone, Liraglutide, Liver fibrosis, Autophagy, Inflammasome, Liver regeneraton

## Abstract

**Background:** Liver fibrosis is a significant complication of chronic liver diseases. While Pirfenidone (PFD) and Liraglutide (LIR) have shown promise individually in treating fibrosis, their combined effect on autophagy and NLRP3 inflammasome pathways remains unexplored.

**Methods and Results:** To investigate the protective effects of combined LIR and PFD therapy on autophagy and NLRP3 inflammasome, fifty male Wistar rats were divided into five groups: Sham, BDL, BDL+PFD (200 mg/kg), BDL+LIR (600 µg/kg), and BDL+PFD+LIR combination. Following 20 days of treatment, liver tissues were analyzed for histological and immunohistochemical (IHC) changes, biochemical parameters, and molecular markers of fibrosis, autophagy, and inflammasome activation. The combination therapy significantly reduced liver damage markers (ALT, AST, ALP), decreased ECM deposition, and improved histological parameters compared to monotherapy. Combined treatment effectively suppressed inflammatory markers (NF-κB, TNF-α) while increasing anti-inflammatory IL-10. Furthermore, the combination therapy modulated autophagy markers (Beclin 1), cathepsib B and reduced NLRP3 inflammasome activation (NLRP3, Caspase 1, IL-1β, IL-18) more effectively than either drug alone. IHC staining of Ki-67 and HepPar-1 showed that combination therapy increased liver regeneration.

**Conclusions:** PFD and LIR combination therapy demonstrates superior therapeutic efficacy in treating BDL-induced LF through elevate liver regeneration and modulation of autophagy and NLRP3 inflammasome pathways, suggesting a promising treatment strategy for LF.

## Introduction

The gradual state of human cholestatic liver disease is followed by cirrhosis, fibrosis, and liver failure [1]. Hepatic tissue experiences several structural and metabolic alterations in cases of extensive liver injury, such as bile duct ligation (BDL)-induced liver fibrosis (LF), which is a complex pathophysiological process. Hepatic stellate cells (HSCs) are activated, which leads to the creation of fibrous tissue and an increase in collagen production in the extracellular matrix (ECM) [2]. By releasing cytokines that cause inflammation and forming fibroblasts in response to different traumas, hepatocytes contribute to the pathophysiology of LF.

Autophagy is a lysosomal-mediated, evolutionarily maintained cellular mechanism that breaks down damaged organelles, protein aggregates, and other macromolecules in the cytoplasm. It controls cell death in both healthy and diseased settings [3, 4]. Liver disorders, such as hepatic fibrosis, are associated with autophagy [5]. While studies have demonstrated autophagy’s control over LF, its precise role in this condition remains unclear. According to earlier research, autophagy controls the expression of genes linked to LF, such as the transforming growth factor TGF-β [6, 7]. There is mounting study that the NLRP3 inflammasome signaling pathway in LF is regulated by autophagy [8].

One of the NOD-like receptors (NLRs) involved in the innate immune response is NLRP3 (NOD-like receptor, pyrin domain-containing 3) [9]. The inflammasome, a protein complex formed by NLRP3, triggers caspase-1 activation, which leads to the maturation and release of pro-inflammatory cytokines such as IL-1β and IL-18 [10]. Therefore, in response to pathogen or damage-associated molecular patterns by both microbial and nonmicrobial stimuli, the NLRP3 inflammasome signaling pathway controls a number of host innate immune defense pathways [11, 12]. It has been demonstrated in research on fibrosis (lung, kidney, etc.) that mice lacking NLRP3 exhibit reduced cytokine and interleukin output. Every piece of data suggests that NLRP3 is a key regulator of fibrosis [14 ,13]. Previous research on LF showed that NLRP3 deficiency reduced inflammation, fibrosis, and liver damage [8].

Pirfenidone (PFD, 5-methyl-1-phenyl-2-(1 H)-pyridone) was authorized via the U.S. Food and Drug Administration in 2014 to treat idiopathic pulmonary fibrosis. By lowering pro-fibrotic cytokines (such as TGF-β1), PFD shown an anti-fibrotic impact. These cytokines slow the progression of fibrosis through a number of methods. These processes include fibroblast proliferation, collagen deposition, epithelial-mesenchymal transition (EMT), and attenuating inflammatory markers [15–17]. Therefore, there have been continuous attempts to investigate a novel therapeutic approach or an alternate treatment for LF. PFD has not yet received approval for treating human LF , however clinical trials are underway [18]. Although there is yet no known molecular target, PFD has been shown to prevent HSC activation and fibrosis in rodents [19–22]. Additionally, it was proposed that PFD might target hepatocytes and other types of hepatic cells [23].

Combination drug therapy appears to be a promising treatment strategy for this high-mortality condition. Since it is difficult to get adequate outcomes with single-drug treatment, multi-drug combination therapies have gained more attention [24]. Because of its capacity to stabilize liver function in hepatic illness, Ligarglutide (LIR), a glucagon-like peptide-1 receptor (GLP-1R) agonist, may be a viable choice for combination therapy. The primary approved use of LIR, an insulinotrophic hormone, is in the management of type 2 diabetes [25]. Many tissues have high levels of GLP-1 expression, with the liver having the highest levels. Both in healthy and diseased situations, GLP-1R activity is crucial for liver function. According to current data, LIR improves the hepatic histological aspects of NAFLD [26].

Furthermore, it has been demonstrated to have a favorable effect on the cellular autophagic response during liver dysfunction and to have very minimal impacts on inflammation brought on by lipotoxicity. In rats with type 2 diabetes, LIR can trigger the cellular autophagic response [27, 28]. LIR is becoming a more appealing therapy option for LF in light of this body of research. Amazingly, we demonstrated in our prior in vivo work that PFD therapy had strong impacts on the regeneration of LF. Further proof of the increased effectiveness of PFD and LIR combination therapy on liver regeneration is therefore valuable. Therefore, we postulated that autophagy might control LF following BDL by controlling the NLRP3 inflammasome. Thus, the purpose of this work was to examine the protective effects of combined LIR and PFD therapy on autophagy and the NLRP3 inflammasome in wistar rats with BDL-induced LF.

## Materials and Methods

### ✓ Animal

Fifty adult male wistar rats weighing 250 ± 300 g were acquired from the Pasteur Institute of Iran’s Laboratory Animal Center. They were housed in a 12-hour light and dark cycle with adequate ventilation in the precise measured ambient conditions of 25±2 °C and 50% humidity. They had unlimited access to tap water and a typical diet of rodent pellets. The Endocrinology and Metabolism Research Institute’s Ethical Committee at Tehran University of Medical Sciences gave its approval to the study (IR.TUMS.AEC.1402.137). All procedures involving animals were conducted in accordance with the guidelines of the Tehran University of Medical Sciences’ Animal Care and Handling Committee, following relevant regulations and the ARRIVE guidelines for ethical animal research.

### ✓ Experimental design

Before the allocation of the rats, body weight was measured. As presented in Fig. 1, animals were grouped into five. Sham groups (n = 10), the BDL group (model group, n=10), and treatment group including: the BDL+ PFD (200 mg/kg body weight, n=10) group, the BDL+LIR (600 µg/kg body weight, n=10) group, and the BDL+ PFD + LIR (PFD=200 and LIR= 600 µg/kg body weight, n=10) group. PFD and LIR doses were selected based on previous research [29, 30]. Prior to biliary obstruction (BDL model induction), rats were given ketamine-xylazine intraperitoneally (i.p.) to induce anesthesia. as previously described [31]. The sham operation just underwent the same laparotomy, but the bile duct was not ligated. After a 5-day acclimation period, all the treatments were administered via gavage once per day for 20 days. An equivalent volume of physiological saline was given to the animals in the model and sham groups.

**Figure 1.**
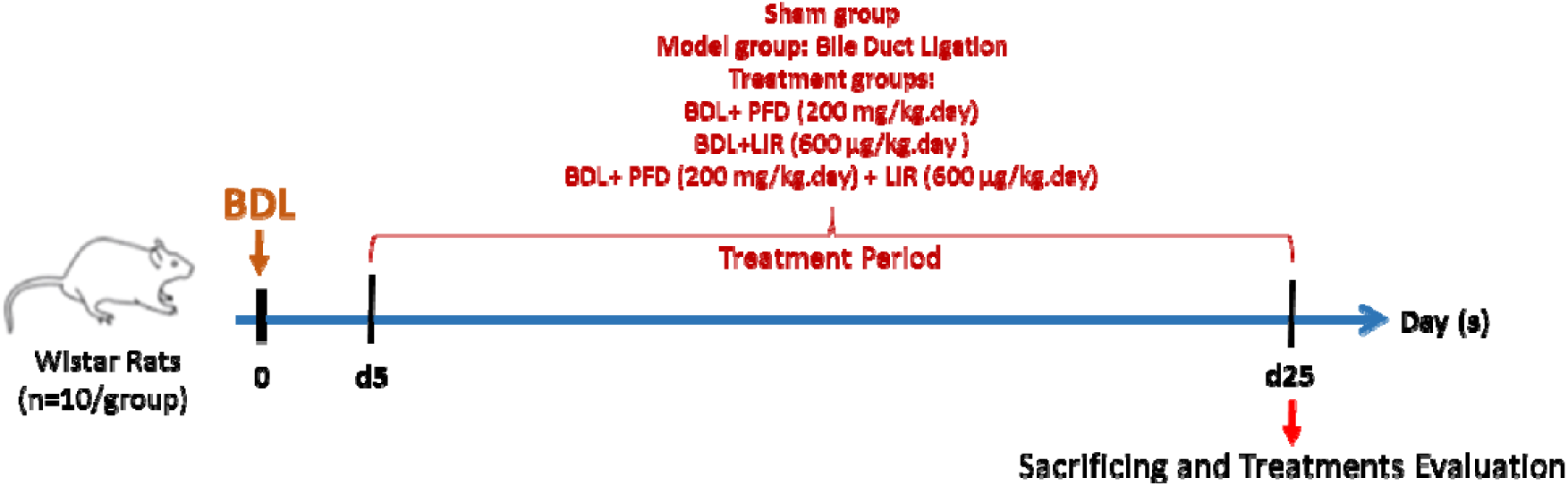
The experimental design and timeline for the study involved BDL-induced liver fibrosis (LF) and corresponding treatment protocols across different rat groups. Liver fibrosis was induced via bile duct ligation (BDL) on day 0. Treatments with PFD, LIR, or their combination commenced five days after BDL induction and were administered daily for 20 consecutive days. Animals were euthanized on day 20 for subsequent analysis.

### ✓ Collection of tissues and blood sample

The rats were sacrificed at the end of the experiment by intraperitoneal (i.p.) injection of a high-dose ketamine/xylazine cocktail, and a heart puncture was used to collect blood samples. The livers were taken out right away. At the conclusion of the experiment, body weights were assessed, and the liver index—or liver/body weight ratio—was estimated. After centrifuging blood samples for 10 minutes at 3000 rpm, a serum specimen was obtained and kept at -80 °C for additional examination. For molecular investigations, hydroxyproline (HYP) content, immunohistochemical, and histological exams, liver tissue was split into many halves.

## Biochemical and molecular studies

### ✓ Biochemical parameter measurement

Alkaline phosphatase (ALP), Aspartate aminotransferase (AST), and Alanine aminotransferase (ALT) levels in serum were assessed using enzymatic methods in compliance with the kit’s instructions (Pars Azmun, Iran).

### ✓ Histological analysis

Haematoxylin and eosin (H&E), Masson trichrome, and Sirius Red were used to stain fixed sections of liver samples. An experienced pathologist blindly evaluated and rated each histopathological sample using the fibrosis scoring system [32].

### ✓ Immunohistochemical analyses

Liver tissue samples from Wistar rats were subjected to immunohistochemical staining for several markers: HepPar-1 (Anti-Hepatocyte Specific Antigen antibody, mouse monoclonal [OCH1E5], GeneTex, USA, diluted 1:100), Alpha smooth muscle actin (α-SMA, Rabbit Anti-ACTA2 Polyclonal antibody, biorbyt, UK, diluted 1:500), and Ki-67, a proliferation indicator (rabbit monoclonal antibody [SP6], BIOCARE MEDICAL, USA, diluted 1:50).

Staining was carried out using a Mouse and Rabbit Specific HRP/DAB Detection IHC kit (Abcam, UK, catalog number ab64264). Following deparaffinization and rehydration, the sections underwent antigen retrieval and were incubated with both primary and secondary antibodies. The stained sections were then analyzed microscopically, and positive cells for each marker were counted in 12 consecutive fields to assess expression percentages, which were subsequently evaluated using Image J software.

### ✓ Hydroxyproline measurement

The hydroxyproline level in liver tissues was measured using commercial hydroxyproline detection kits (Kalazist Life Sciences, Hamedan, Iran) in accordance with the kit’s instructions.

### ✓ RNA isolation and RT-PCR

The frozen liver tissues’ total RNA was extracted by the Trizol reagent (Yektatajhize, Iran). RNA was reverse-transcribed to cDNA using the Revert Aid cDNA Synthesis Kit (Yektatajhiz, Iran). On an ABI-Step One (Applied Biosystems, USA), quantitative real-time PCR was performed using Real Plus 2x Master Mix SYBR Green (Amplicon, Denmark). Primer sequences are available upon request. The results were standardized to the GAPDH levels and then evaluated using the comparative 2−ΔΔCt method for relative quantification.

### ✓ Western blotting

In order to extract proteins, liver tissues were homogenized in lysis buffer. The bicinchoninic acid assay (BCA) kit (Thermo Fisher Scientific, UK) was applied to quantify the protein concentration. The Beclin 1, Cathepsin B (CTSB), NLRP3 and Caspase 1 and GAPDH immunoblotting was carried out. Secondary antibodies were then applied to the blots (Santa Cruz, USA). Finally, the ECL reagent and Image J software (National Institutes of Health, Bethesda, USA) were used to view the immunoblot.

### ✓ Statistical analysis

All data are expressed using the standard deviation (SD) of the mean. For statistical analysis, Prism version 9 (GraphPad Software Inc., San Diego, CA, USA) was used. The data was compared using a one-way analysis of variance (ANOVA). Statistical significance was defined as a value of P < 0.05.

## Results

### LIR and PFD alone and their combination alleviate BDL-induced liver fibrosis in rats

First, we looked into how LIR, PFD, and their combination affected LF in vivo. Rats given BDL revealed pathological liver changes in comparison to normal livers; however, the pathological liver changes were significantly lessened in the LIR, PFD, and LIR+PFD supplementation groups. H&E staining showed that the fibrosis of LIR, PFD and LIR+PFD were markedly lower than the BDL rats, as evidenced by ameliorated hepatic steatosis, necrosis, and fibrotic septa (FIG. 2a). LIR and PFD significantly reduced the liver/body weight ratio, which was significantly elevated by BDL-induction (FIG. 2b). Furthermore, LIR+PFD combination therapy dramatically decreased serum levels of three well-known indicators for hepatocyte injury—ALT, AST, and ALP—when compared to rats given BDL (Fig. 2c). Taken together, LIR+PFD combination therapy is an effective therapy to decrease hepatic injury caused by BDL-induction compared to LIR and PFD monotherapy.

**Figure 2.**
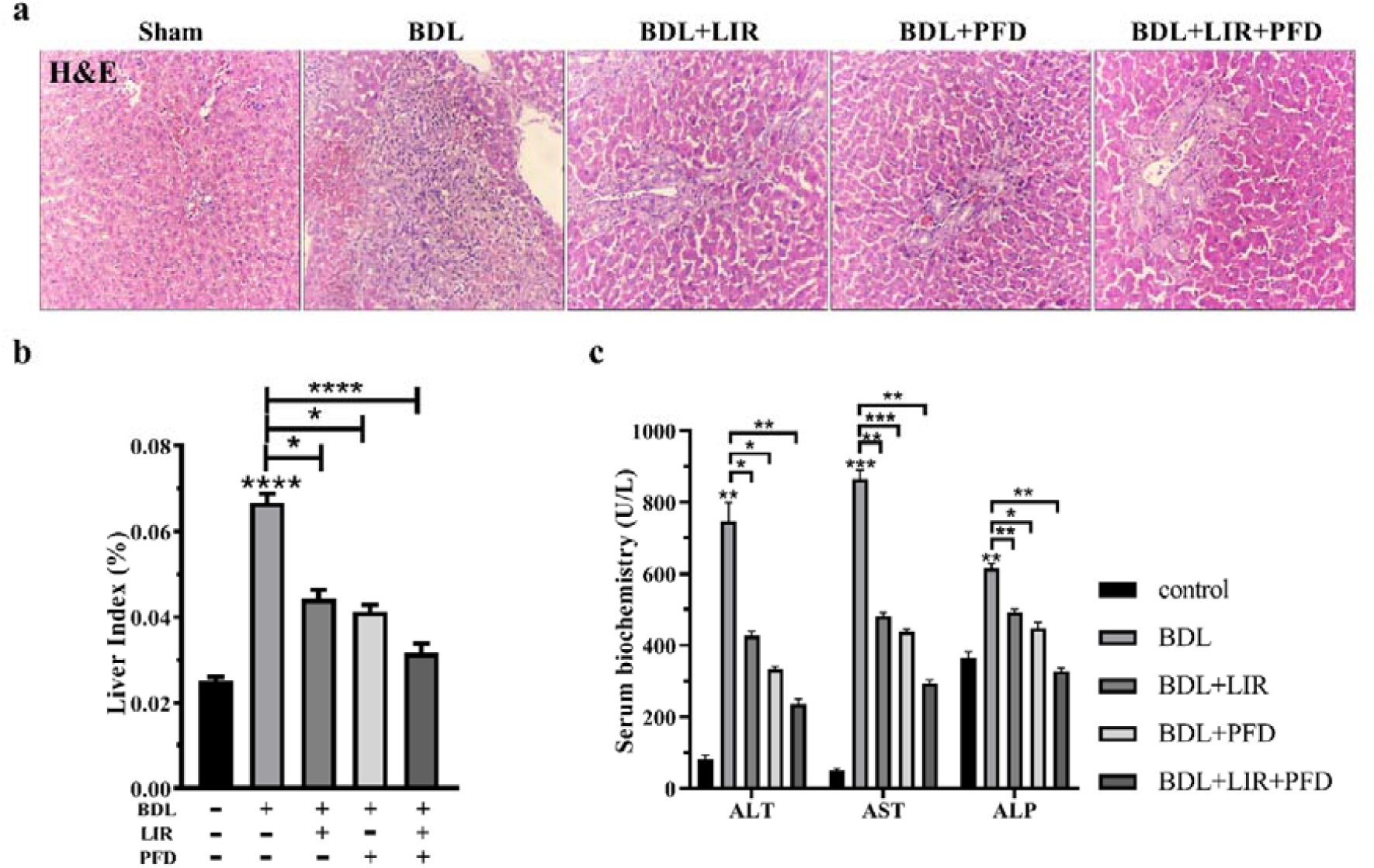
Effect of LIR and PFD alone and combination of LIR+PFD on the BDL-induced rat. (a) H&E staining of different groups for histological examination. Scale bar: 100 µm and magnification: X100. (b) The rat liver/body weight ratio. (c) The serum levels of ALT, AST and ALP.

### LIR and PFD alone and their combination alleviate BDL-induced ECM deposition in rats

The sham rats exhibited typical liver tissue structure as shown by Masson trichrome and Sirius Red staining. The BDL group showed marked proliferation around the bile duct and collagen deposition. While collagen deposition and necrosis surrounding the portal tract were significantly decreased in LIR, PFD and LIR+PFD supplementation groups (Fig. 3a).

**Figure 3.**
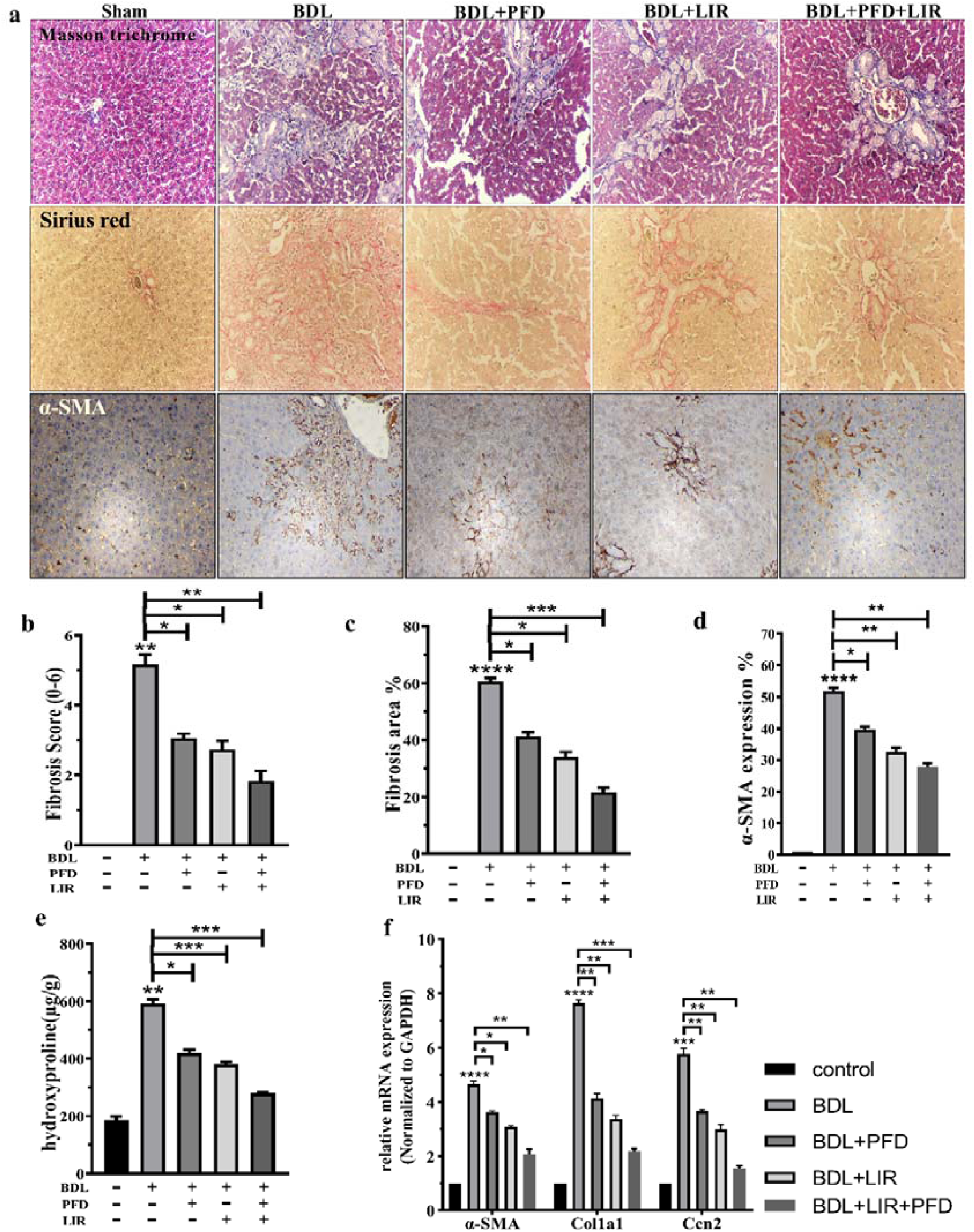
Effects of LIR and PFD alone and LIR+PFD combination on the BDL-induced ECM deposition in rats. (a) Masson trichrome and Sirius Red staining for histological examination and Sections of liver tissue immunostained against α-SMA for different groups. Scale bar: 100 µm and magnification: X100. (b) fibrosis scores and (c) fibrosis area; (d) The α-SMA expression (%) was recorded by counting the positive cells; (e) liver tissue HYP content; and (f) the mRNA expression level of α-SMA, Col1a1, and Ccn2.

Although the fibrosis score and area were dramatically reduced in all treatment groups, the combination (LIR+PFD supplementation) group may be more protective against all pathological liver alterations (Fig. 3b&c). We used immunohistochemical labeling to assess α-SMA expression, aiming to evaluate the effect of PFD and LIR combination therapy on collagen synthesis. In the sham group, minimal α-SMA expression was observed around the bile ducts and nearby blood vessels. Bile duct ligation (BDL)-induced liver fibrosis was confirmed by a significant increase in α-SMA expression. Notably, treatment with the drugs resulted in a marked decrease in α-SMA levels in the BDL groups, with the combination of LIR and PFD showing the most pronounced reduction. These findings suggest that the LIR+PFD therapy effectively diminishes extracellular matrix deposition compared to untreated BDL-induced fibrosis (Fig. 3d). The HYP content in tissues was evaluated as displayed in Fig. 3e. The HYP content increased significantly in the BDL group compared with the sham group. Following all of the treatments, these changes were preserved. In addition, combined drug treatment (LIR+PFD supplementation) was more effective in the reduction of HYP than the single ones compared with the BDL group. Furthermore, to evaluate the supportive effect of PFD and LIR and their combination against ECM deposition in BDL-induced LF, we examined the crucial ECM-producing markers using RT-PCR (Fig. 3f). In the BDL group, a meaningful elevation in α-SMA, Col1a1 and Ccn2 was detected compared to the sham group. As expected, all the treatments significantly decreased the mRNA levels of these genes. The LIR+PFD combination was more effective in the reduction of α-SMA, Col1a1, and Ccn2, which would lessen the accumulation of ECM. It seems that the LIR+PFD combination is an effective treatment of BDL-induced LF.

### Combination therapy of LIR and PFD reduces inflammatory markers in rats with BDL-induced liver fibrosis

In order to assess how well PFD and LIR and their combination reduce inflammatory responses, the mRNA expression of NF-κB, TNF-α and IL-10 were quantified by RT-PCR. The results showed that BDL-induced LF led to a significant elevate in NF-κB and TNF-α and a decrease in IL-10 mRNA level compared to the sham group. All treatment groups showed a significant decrease in NF-κB and TNF-α and an increase in IL-10. In addition, LIR+PFD combined therapy was more effective in inflammation balance restoration compared with the BDL group (Figs. 4a-c). These findings suggest that LIR+PFD combination therapy reduced hepatic inflammatory infiltration in LF induced by BDL.

**Figure 4.**
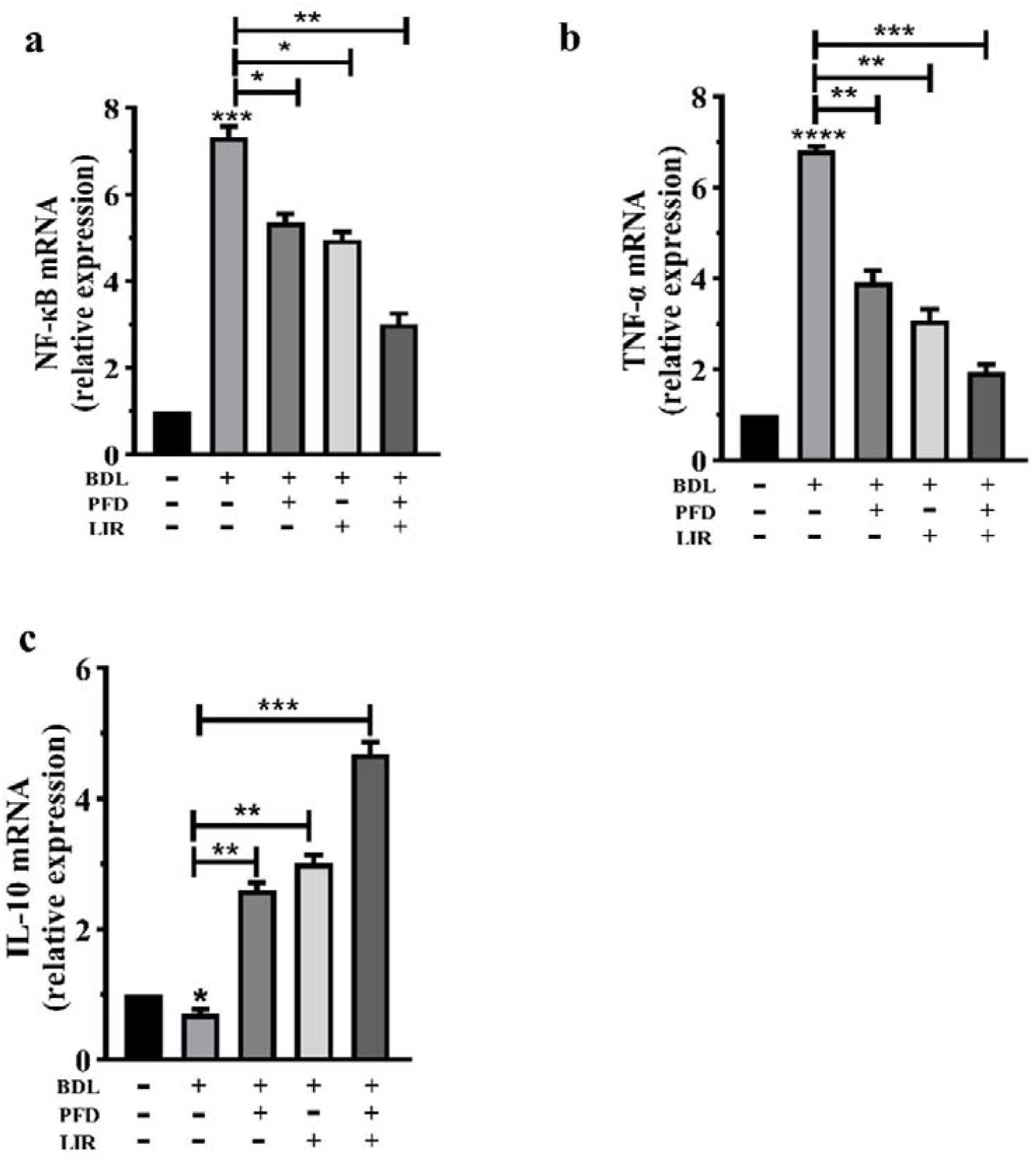
Effects of LIR and PFD alone and LIR+PFD combination on the BDL-induced inflammatory response in rats. The mRNA levels of (a) NF-κB, (b) TNF-α, and (c) IL-10.

### Combination therapy of LIR and PFD alleviates autophagy and NLRP3 inflammasome activation in rats with BDL-induced liver fibrosis

To assess the effects of LIR and PFD alone and their combination on autophagy and NLRP3 inflammasome activation, Beclin 1 as an autophagy marker and NLRP3, Caspase 1, Asc, IL-1 β and IL-18 as NLRP3 inflammasome markers, as well as CTSB, were measured using western blot and RT-PCR. The protective effect of LIR and PFD in relation to autophagy activity was assessed by the changes of key autophagy regulatory proteins in BDL-induced LF rats. The BDL group displayed significantly higher protein expression of Beclin-1 compared to the sham group. The treatment of LIR and PFD alone and their combination reduced protein expression of Beclin 1 protein expression. LIR+PFD combined therapy was more effective than LIR and PFD monotherapy. In addition, rats in the BDL group revealed a significant elevation in caspase-1 activity and NLRP3 protein content as well as IL-1β and IL-18 mRNA expression compared with the sham group. In contrast, the caspase-1, NLRP3 protein expression and IL-1β and IL-18 mRNA expression were decreased significantly upon LIR, PFD and LIR+PFD combination administration, with superiority of the combination therapy over the monotherapy when compared with the BDL group. Compared with the sham group, BDL-induced LF significantly increased the CTSB protein content (Figs. 5a-e). Treatment with LIR, PFD and LIR+PFD combination, in contrast, significantly decreased the level of CTSB protein. The LIR+PFD combination was superior to each drug alone.

**Figure 5.**
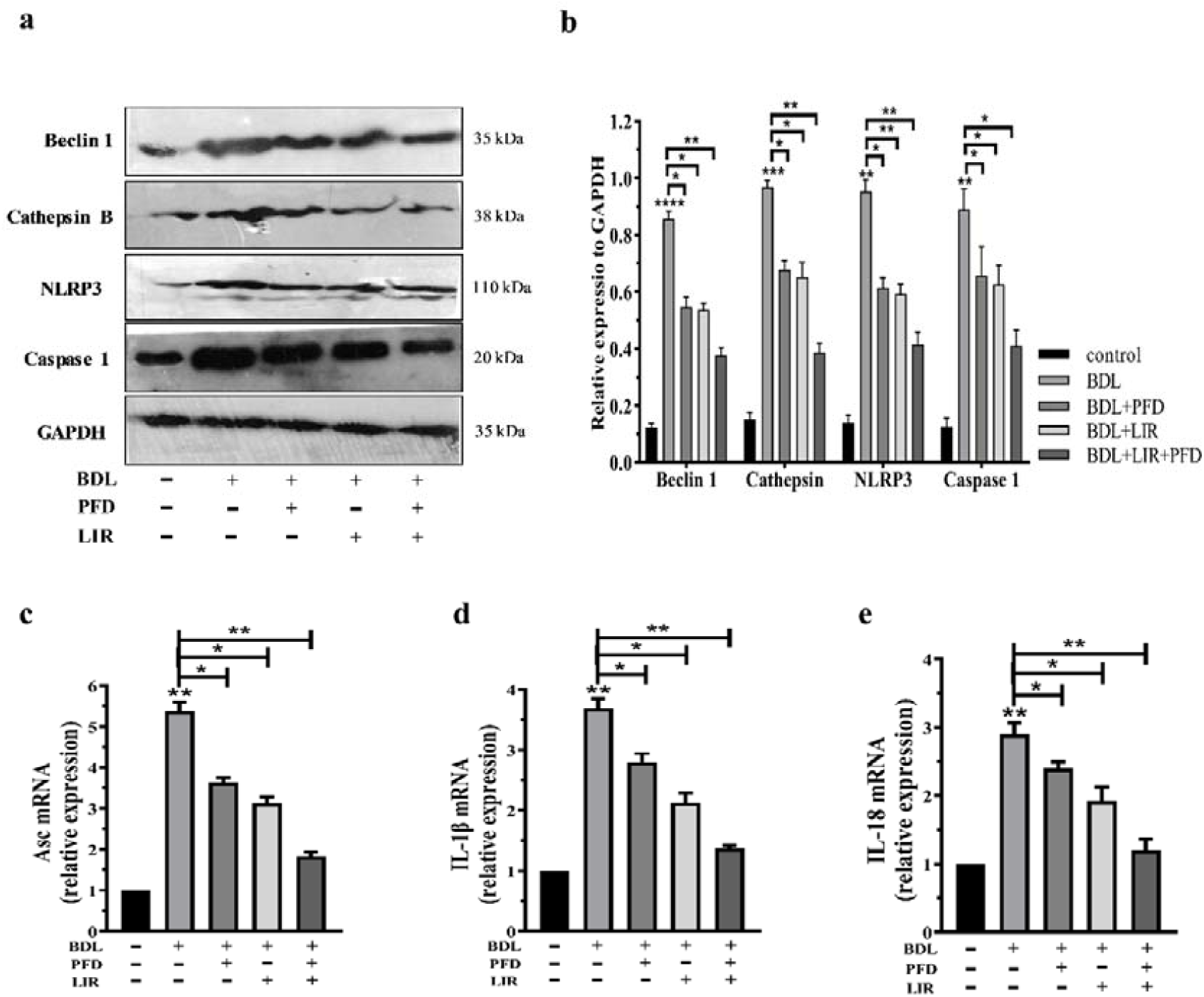
Effect of LIR and PFD alone and combination of LIR+PFD on the elevated levels of autophagy and NLRP3 inflammasome activation in the BDL-induced LF rats. (a) Representative protein expression of Beclin 1, Cathepsin B, NLRP3 and Caspase 1 in liver tissue using western blotting. The data were normalized to GAPDH levels. (b) Densitometric quantitation of Beclin 1, Cathepsin B, NLRP3 and Caspase 1. The mRNA levels of (c) Asc, (d) IL-1β and (e) IL-18.

### Combination therapy of LIR and PFD promotes liver regeneration in rats with BDL-induced liver fibrosis

To assess the effects of PFD, LIR, and their combination on HepPar-1 and Ki-67 expression, immunohistochemistry was performed on liver tissues from the experimental groups. Representative images of HepPar-1 and Ki-67 staining are shown in Figure 6a. Liver sections from the control animals exhibited a characteristic dense, granular cytoplasmic staining pattern. In contrast, the BDL group showed a weakened and heterogenous staining across hepatocytes, indicating that bile duct ligation negatively impacted HepPar-1 expression. Hepatocytes in the PFD and LIR groups demonstrated an intermediate level of staining intensity. Notably, liver sections from rats receiving combination therapy displayed strong, dense, granular HepPar-1 immunostaining (Fig. 6b). Additionally, histomorphometric analysis of Ki-67 immunostaining revealed a mild increase in nuclear Ki-67 positivity in BDL sections compared to controls. Treatment with LIR+PFD significantly elevated Ki-67 expression relative to the BDL group, indicating potential regenerative effects (Fig. 6c).

**Figure 6.**
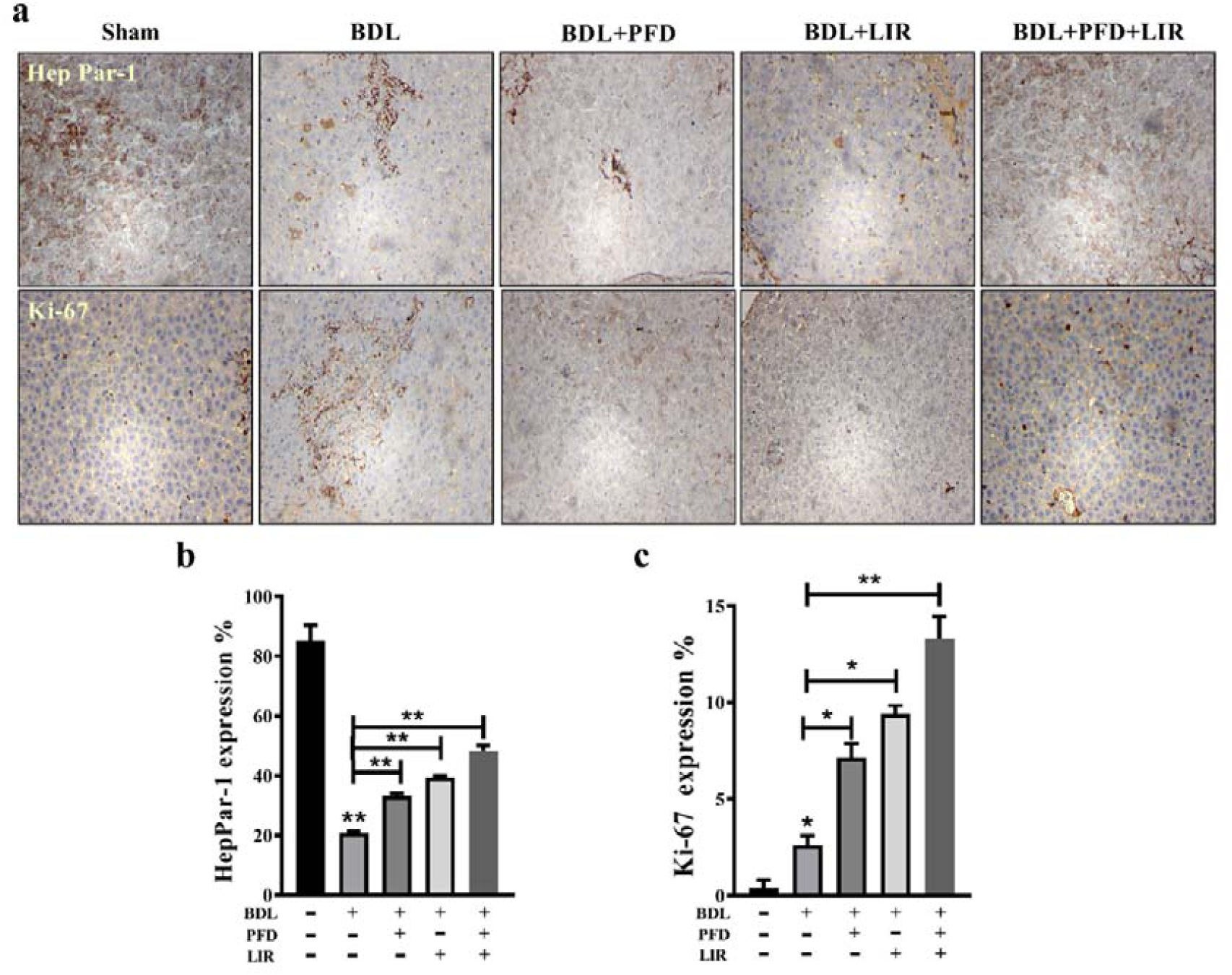
The effects of LIR, PFD, and their combination on HepPar-1 and Ki-67 expression were evaluated in liver sections from BDL-induced wistar rats using immunohistochemistry. (a) Representative images of liver tissue stained for HepPar-1 and Ki-67 are shown, with a scale bar of 100 µm. Quantitative analysis involved counting the number of positive cells, and the results are expressed as the percentage of HepPar-1 and Ki-67 positive cells in each group (b and c).

## Discussion

Hepatic fibrosis is the last typical mechanism for almost all kinds of chronic liver disease [1]. The bile duct epithelium is the primary site of most pathological abnormalities in BDL-induced LF, which is its distinguishing characteristic. It has a high rate of morbidity and a poor prognosis. Much progress has been made in understanding the biology of LF and the treatments that are currently accessible [33, 34]. Specifically, we emphasized autophagy, inflammation, and fibrogenesis as important barriers to fibrosis treatment, all of which are present in the current model of BDL-induced LF. Given the present obstacles in selecting the best drugs for the treatment of LF, combination therapy may be seen as a reasonable alternative approach [35, 36]. The goal of the current study was to determine if PFD and LIR combinations may alleviate hepatic steatosis in the model of rats with BDL-induced LF. This is the only study that shows the combination of PFD and LIR can help decrease LF.

Because PFD is an anti-fibrotic substance, it is applied to treat lung fibrosis. There is great potential for this drug to effectively treat LF [37], as our recent research has also demonstrated [32]. Thus, the demand to create more effective treatment plans for people with LF is increasing [38]. LF responds well to PFD combined therapy [39]. In order to learn more about combination therapy, we also worked on this concept. When GLP-1 receptor LIR is present, human hepatocytes internalize the GLP-1 receptor and lessen liver damage [40, 41]. LIR reduced the expression of NLRP3 in the liver of mice and improved hepatic steatosis [42]. In a model of diabetic retinopathy, GLP-1 decreased levels of pro-inflammatory cytokines and NLRP3 [43]. More findings demonstrated that LIR enhanced the expression of proteins linked to autophagy in rats in a dose-dependent manner. These findings imply that LIR reduces liver damage by triggering autophagy [44].

We investigated whether the combination of PFD and LIR protected from the progression of liver injury in a rat model of common BDL. The hepatoprotective efficacy of PFD+LIR combination therapy has been assessed using animal models of BDL-induced LF. PFD+LIR significantly treated liver damage caused by BDL. According to liver enzyme testing, liver index, and H&E staining, PFD+LIR significantly decreased liver damage. PFD and LIR in combination cause a more noticeable improvement in BDL-induced LF, according to analysis of the liver’s HYP concentration, Masson trichrome, and Sirius Red staining. It appears that reduced liver index, HYP concentration, enhanced HepPar-1 immunostaining and ultimately collagen deposition may be partially responsible for the positive benefits of combined PFD and LIR treatment on LF. In the BDL fibrosis model, PFD has been demonstrated to have an anti-fibrotic effect [29]. Furthermore, LF was reduced by LIR due to its strong anti-inflammatory, hepatic microvascular, and anti-fibrotic properties [45]. The potential of PFD and LIR to support liver regeneration was also demonstrated by our investigation. Increased HepPar1 and Ki-67 immunostaining, indicators of functional hepatocytes, coincided with these alterations [46]. Consistent with earlier research, our data show that PFD and LIR dramatically reduce transaminases, improve the liver index, and improve HepPar-1 immunostaining, thereby mitigating BDL-induced liver injury [47–49]. All of these outcomes point to better liver function. We also noticed improvements in the hepatic architecture in addition to these improvements. PFD treatment produced high, diffuse nuclear Ki-67 immunostaining, but BDL-group rats showed modest Ki-67 immunostaining, indicating hepatocyte proliferation as part of the adaptive response to liver tissue damage [50–52]. This finding implies that PFD and LIR might cause a more noticeable rise in hepatocyte proliferation, which could result in better liver regeneration.

In the present study, we showed that in fibrotic tissues, PFD+LIR decreased the expression of collagen deposition marker mRNA. These results imply that PFD+LIR antifibrotic activity may be mediated by molecular pathways such as α-SMA, Col1a1, and Ccn2 regulating ECM deposition.

According to earlier research, inflammasomes are responsible for the acute inflammation that results from BDL-induced liver damage. It has been determined that NLRP3 is a crucial modulator of liver inflammation [53]. Increased caspase-1 expression, IL-18, and the cleavage of pro-IL-1β to IL-1β are all brought on by the activation of NLRP3 inflammasomes. When IL-1β interacts with the IL-1 receptor (IL-1R), IL-1R-expressing cells are recruited into the liver, which intensifies the inflammatory response [54]. Thus, by lowering inflammation, inflammasome inhibition may help mitigate liver damage brought on by BDL. Our findings showed that rats with BDL-induced liver injury had considerably higher mRNA levels of Asc and protein expressions of NLRP3 and caspase-1, which is in line with earlier research. In line with our data According to YU et al., LIR reduces liver disease via blocking the NLRP3 inflammasome [55]. Furthermore, PFD has been shown to reduce lung fibrosis and inflammation by preventing the activation of the NLRP3 inflammasome [56]. Other studies have documented NLRP3 inflammatory expression in a rat model of LF brought on by common BDL [57]. The significance of NLRP3-mediated inflammation in the pathophysiology of BDL-induced liver damage was validated by our findings. Rats treated with PFD and LIR had lower levels of NLRP3 expression and IL-18 cytokine production. It’s interesting to note that model groups showed a protective effect from PFD+LIR combination therapy, in contrast to BDL rats. These findings showed that the suppression of the NLRP3 inflammasome was linked to PFD+LIR’s hepatoprotective effects against BDL-induced LF.

Autophagy is one self-digesting mechanism to preserve cellular homeostasis [58]. The autophagy process addresses both fused lysosomes, which produce autolysosomes, and built autophagosomes. The misfolded proteins are then absorbed, and the injured organelles are broken down [59]. Recent data suggested that autophagy might be a crucial defense against liver damage brought on by BDL. In addition to removing damaged mitochondria, autophagy can supply the energy needed for the synthesis of adenosine triphosphate (ATP) [60]. Acute hepatic damage may therefore be lessened by therapies that increase the liver’s autophagy flux [58]. According to the current study, BDL surgery raised the protein beclin-1, indicating a higher flow of autophagy in the liver tissues. Mechanistically, BDL is linked to increased bile duct proliferation and the production of ROS in the mitochondria, which in turn trigger autophagy. It’s interesting to note that our research showed that PFD+LIR combination treatment greatly decreased autophagy activity due to much lower beclin-1. It is unclear exactly how the PFD+LIR combination medication lowers autophagy, but one theory is that it does so by decreasing autophagy, which in turn lowers hepatocyte inflammation and necrosis.

The link between NLRP3 suppression and autophagy in response to the PFD+LIR combination was also investigated in the current study. More research indicates that NLRP3 and autophagy seem to be regulated by one another [61].

According to the earlier research, macrophage autophagy alters the activation of the inflammasome, which in turn alters the liver’s immunological reactivity [62]. Additionally, autophagy suppression leads to aberrant apoptosis induction, which worsens hepatocyte dysfunction. [63]. Functional autophagy shields the liver against harm, according to all of the data above. The accumulation of depolarized mitochondria brought on by autophagy suppression results in the release of molecules that activate endogenous inflammasomes, proving that autophagy is necessary for the destruction of NLRP3 [64, 65]. Autophagy limits the development and generation of IL-1β and IL-18, decreases caspase-1 activation, and encourages the destruction of the NLRP3 inflammasome [66].

The lysosomal cysteine protease CTSB has been implicated in the development of autophagy-induced inflammasomes [67, 68]. Here, we report that the elevated level of CTSB in the BDL group due to liver injury may have served as a stimulus that activated the NLRP3 inflammasome, promoted the secretion of caspase-1-dependent proinflammatory cytokines and other inflammasome-related factors, and may have accelerated the pathogenesis of LF.

Furthermore, the combination of PFD and LIR decreased the levels of liver CTSB brought on by BDL. Additional study by other scientists has shown that NLRP3 can be activated by the release of CTSB following injury [69]. In line with our findings, excessive autophagy activation increases lysosomal instability and is linked to NLRP3 activation brought on by liver damage, both of which are likely contributing factors to the inflammation that encourages LF. According to these findings, which are consistent with our investigation, lowering CTSB could prevent its release, hence reducing the inflammatory response [70].

According to recent research, autophagic mechanisms and inflammation interact in a complicated way. BDL-dependent inflammasome activation is made possible by autophagy medication therapy, indicating that autophagy often inhibits the activation of inflammasomes caused by liver damage. Autophagosomes may target inflammasomes for destruction, and reactive oxygen species may be a potential target that indirectly inhibits inflammasome activity [71]. Our findings, however, point to a potential interaction between the proinflammatory cytokine production and the adverse consequences of excessive autophagy activation. Autophagy reduces the formation of CTSB after PFD+LIR combination, which can alter intracellular inflammasomes. This, in turn, activates NLRP3 and inflammatory caspases. The findings presented here are also in line with recent research showing that autophagy and NLRP3 inflammasomes may be essential for reducing liver damage brought on by BDL [57, 72–74].

## Conclusion

The current study offers convincing evidence that PFD and LIR together may enhances regenerative response and improve autophagy and the NLRP3 inflammasome mechanism, which may help to alleviate LF (Fig. 7). Our results offer novel perspectives on the mechanisms behind the release of proinflammatory cytokines linked to autophagy, demonstrating that autophagy is also associated with the release of NLRP3 inflammasomes. Examining the intricate relationship between autophagy and the NLRP3 inflammasome response throughout the course of LF in hepatocytes from hepatic fibrosis and animal models would be intriguing in light of this potential.

**Figure 7.**
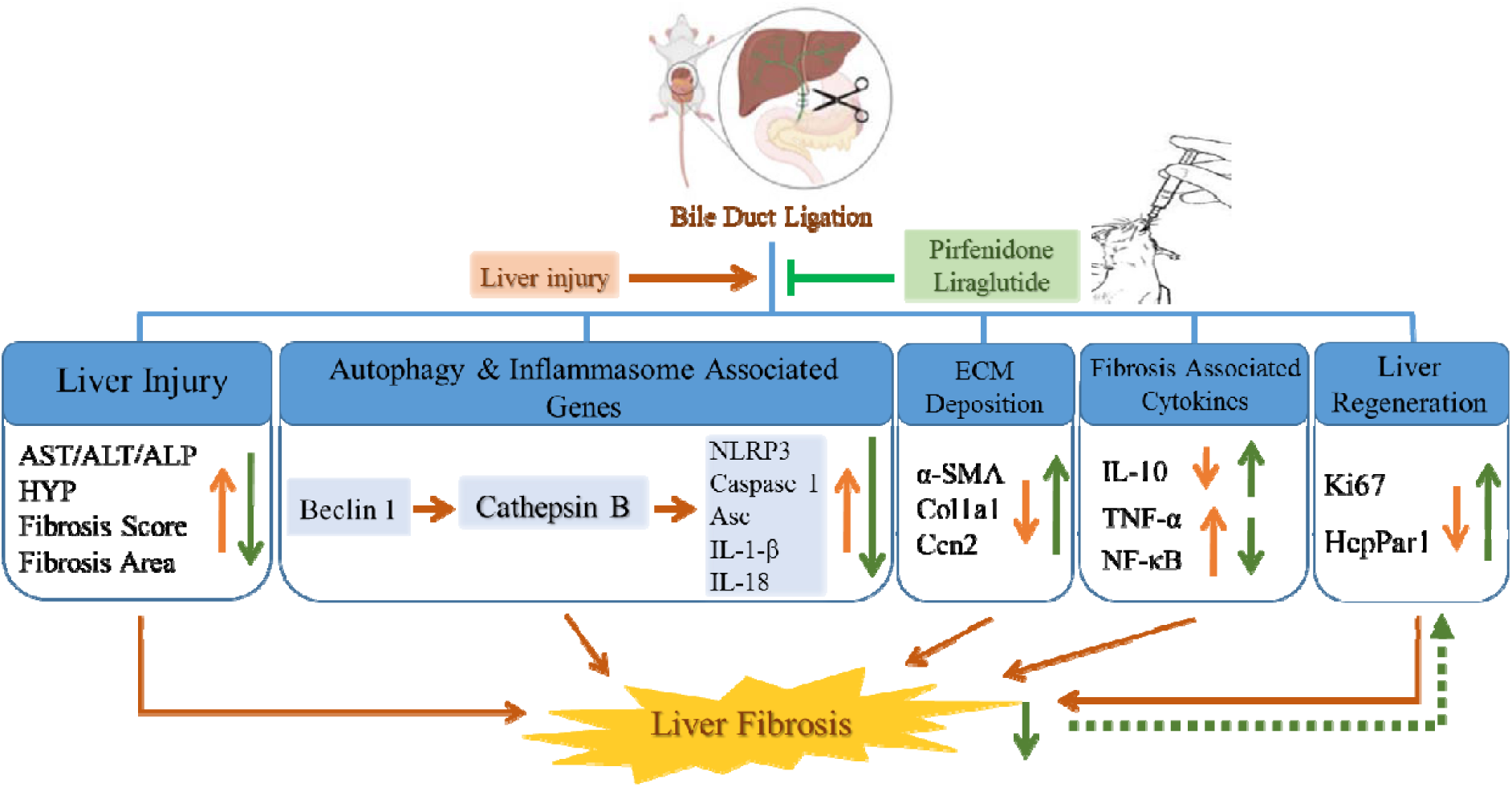
Theoretical schematic of LIR and PFD combination therapy in rats with BDL-induced liver fibrosis.

## Authors’ Contributions

ZY: Investigation, data collection, writing the original draft and editing, formal analysis, and methodology. RR: Writing, review, editing, and methodology. SK: formal analysis, review, editing. ASL: Supervision, investigation, editing, validation, visualization, and project administration. MN: Supervision, validation, visualization, and funding acquisition.

## Ethics approval

The Endocrinology and Metabolism Research Institute’s Ethical Committee at Tehran University of Medical Sciences gave its approval to the study (IR.TUMS.AEC.1402.137).

## Acknowledgement

The authors would like to express their gratitude to the Endocrinology and Metabolism Research Institute, Tehran University of Medical Sciences, for their support of this research.

## Conflict of Interest

The authors declare no conflict of interest, financial or otherwise.

**Abbreviations**
(LF): Liver fibrosis
(PFD): Pirfenidone
(LIR): Liraglutide
(BDL): Bile Duct Ligation
(ALT): Alanine aminotransferase
(AST): Aspartate aminotransferase
(ALP): Alkaline phosphatase
(ECM): Extracellular Matrix
(H&E): Haematoxylin and Eosin
(HSC): Hepatic Stellate Cell
(α-SMA): α-smooth muscle actin
(MFB): Myofibroblast
(TGF-β): Transforming Growth Factor β
(ROS): Reactive Oxygen Species
(HepPar-1): Hepatocyte Paraffin 1
(Hyp): Hydroxyproline
(TNF-α): Tumor Necrosis Factor-alpha
(GAPDH): Glyceraldehyde 3-phosphate dehydrogenase
(NF-κB): Nuclear Factor Kappa B
(NLRP3): NOD-, LRR- and pyrin domain-containing protein 3
(CTSB): Cathepsin B

